# Differences in proteome response to cold acclimation in *Zoysia japonica* cultivars with different levels of freeze tolerance

**DOI:** 10.1101/581488

**Authors:** J.M. Brown, H. McCamy P. Holloway, Michelle DaCosta, Rachael P. Bernstein, Jefferson Lu, Tan D. Tuong, Aaron J. Patton, Jeffrey C. Dunne, Consuelo Arellano, David P. Livingston, Susana R. Milla-Lewis

## Abstract

Zoysiagrasses (*Zoysia* spp.) are warm season turfgrasses primarily grown in the southern and transition zones of the United States. An understanding of the physiological changes that zoysiagrasses undergo during cold acclimation may shed light on physiological phenotypic traits useful in selection of freeze tolerant genotypes. We investigated the relationship between cold acclimation, protein expression, and freeze tolerance in cold-acclimated (CA) and non-acclimated (NCA) plants of *Zoysia japonica* Steud. cultivars ‘Meyer’ (freeze-tolerant) and ‘Victoria’ (freeze-susceptible). Freeze tolerance was assessed using chambers reaching −6, −8, −10, and −12°C. Additionally, meristematic tissues from the grass crowns of ‘Meyer’ and ‘Victoria’ were harvested for proteomic analysis after a four week cold acclimation period. Freeze testing indicated that cold acclimation accounted for a 1.9-fold increase in plant survival compared to the non-acclimation treatment. Overall, proteomic analysis identified 62 protein spots having at least a twofold change in abundance under cold acclimation. Nine and 22 unique protein spots were identified for Meyer and Victoria, respectively, with increased abundance (up-regulated) or decreased abundance (down-regulated). In addition, 23 shared protein spots were found among the two cultivars having differential expression in response to cold acclimation. In Meyer, protein response to cold acclimation was primarily upregulated, while in Victoria, protein response was primarily downregulated. These cold acclimation responsive proteins were found to be involved primarily in transcription, metabolism, protein destination and storage, and energy production. As identified through MALDI-TOF/TOF mass spectrometry followed matching of protein homologues against the NCBI Arabidopsis database, major proteins of interest for their association with cold acclimation were LEA 3, MAPK, SOD, GAST1, Phytochrome A, ATP synthase, AGP, PLD, and PSII. Further investigation of these proteins and their functional categories may contribute to increase our understanding of the differences in freezing tolerance among zoysiagrass germplasm.

## INTRODUCTION

Zoysiagrasses (*Zoysia* spp.) are warm season perennial turfgrasses primarily grown in the warm-humid and transitional climatic zones of the United States. They are geographically constrained to these regions because of their limited winter hardiness. Zoysiagrasses are popular for their low maintenance requirements as well as their slow growth habit, high shoot density, and tolerance to many abiotic stresses such as drought and shade [1]. These attributes make zoysiagrass especially well-suited for home lawns, landscapes, and golf courses and creates a demand for more freeze tolerant cultivars that can be used in the colder climates of the Northern US.

Varying levels of winter injury have been observed among zoysiagrass genotypes. The two most popular species of zoysiagrass in the U.S., *Zoysia japonica* Steud. and *Zoysia matrella* (L.) Merr., have significantly different levels of winter injury and spring green-up as first reported by Forbes and Ferguson [2]. Both in the field and freeze chamber studies, *Z. japonica* genotypes show more average freeze tolerance than *Z. matrella* genotypes [3, 4, 5]. Patton and Reicher [5] reported that the LT_50_s for *Z. japonica* commercial cultivars range from −9.5°C to −11.5°C for ‘Victoria’ and ‘Meyer’, respectively. Additionally, LT_50_s have been found to differ depending on the origin of the tested material and freeze testing protocol [3, 4, 5, 6].

Cold acclimation refers to the natural physiological process that takes place in a plant when it is exposed to low but above freezing temperatures. Studies in zoysiagrass [4, 5, 6], as well as saltgrass (*Distichlis spicata* (L.) Greene) [7], buffalograss (*Bouteloua dactyloides* (Nutt.) Engelm.) [8], bermudagrass (*Cynodon dactylon* (L.) Pers. *× C. transvaalensis* Burt-Davy) [9, 10] and St. Augustinegrass (*Stenotaphrum secundatum* (Walt.) Kuntze) [11, 12, 13] have indicated that warm-season grasses respond well to cold acclimation suffering less injury from freezing when acclimated. Hinton et al., [4] found that across nine different zoysiagrass cultivars with varying LT_50_s, samples collected in the winter (after cold acclimation) were consistently more freeze tolerant in controlled environment freezing tests than samples collected in the spring (non-acclimated/de-acclimated).

Although field evaluation of freeze tolerance provides the most realistic indication of how a cultivar will perform through winter, variable environmental conditions across year and location can make consistent and reproducible winter stress very difficult to attain [14]. Controlled environment freeze tests may be a more cost-effective and efficient way to assess cold acclimation and freeze tolerance of turfgrass species and generally corresponded well with field screenings [8, 14, 15]. This constancy demonstrates that plants most likely undergo similar physiological changes during cold acclimation, freezing, and deacclimation in temperature controlled chambers as in the field. In zoysiagrass, freezing chambers have been successfully used to evaluate the low temperature tolerance of a variety of genotypes [3, 4, 5, 16].

Our understanding of the physiological changes that zoysiagrass undergoes during cold acclimation has been somewhat limited. Zhang et al. [17], investigated response to cold acclimation in zoysiagrass cultivars ‘Meyer’ and ‘Cavalier’ to identify changes in hormones and dehydrin expression. Most notably, they found an increase in abscisic acid and dehydrin levels, and a decrease in cytokinin levels in Meyer compared to Cavalier, which may play a role in winter survival and freeze tolerance. Patton et al. [18, 19] compared levels of compounds suspected to be associated with freeze tolerance such as carbohydrates, organic acids, and soluble proteins at various points in the cold acclimation process. The study found that carbohydrates such as glucose and sucrose increase in stolons during seasonal temperature decreases in zoysiagrass, similar to what is known to occur in other warm season turfgrasses such as saltgrass (*Distichlis spicata* var. stricta (L.) Greene) [7], centipedegrass (*Eremochloa ophiuroides* (Munro) Hack.) [20] and buffalograss (*Buchloe dactyloides* (Nutt.) Englem.) [21]. However, a comparison between LT_50_ and sucrose concentration in CA grasses revealed no relationship with freezing tolerance. Despite this result, evidence suggested that genotypes with poor freezing tolerance produce more sucrose during acclimation than genotypes with high freeze tolerance, and that total reducing sugars such as glucose likely play a role in total freezing tolerance. Proline concentration is known to increase dramatically during cold acclimation due to an uptake in synthesis with freeze tolerance improving with increased proline [19]. Santarius [22] found that proline protects membranes using hydrophobic interactions, osmotic adjustment, and decreased water potential, but it is less efficient as temperatures decline. Lastly, Zhang et al. [6] evaluated changes in lipid composition during cold acclimation, but found no consistent relationship between lipid classes and zoysiagrass freezing tolerance.

Comprehensive studies of the proteins involved in cold acclimation as well as their functions may lead to more accurate and efficient selection of associated genes in zoysiagrass. One such study, by Xuan et al. [23] performed quantitative analysis of protein accumulation in stolon tissues collected from two species, *Z. japonica* cv. Meyer and *Z. matrella* cv. Diamond, during cold acclimation. Forty-five proteins with a two-fold change in expression were functionally identified as playing a role in key functional protein classes such as redox homeostasis, signal transduction, photosynthesis, and energy metabolism among others. A close examination of the proteins involved suggests that a greater energy supply is needed to respond to cold stress in zoysiagrass stolons. Meyer’s ability to respond to cold stress most likely is related to its increased ROS scavenging ability, photosynthesis, protein synthesis, proteolysis, and its greater energy reserves. However, it may be relevant to note that the LT_50_s of the two cultivars in this study was determined through electrolyte leakage, a method which tends to overestimate the LT_50_, rather than whole plant survival. Furthermore, their use of stolons rather than meristematic tissues from the grass crown may lead to different inferences as a result of the specialization of plant tissues. To identify genomic regions which play a role in freezing tolerance, Holloway et al. [24] generated a high density linkage map of *Zoysia japonica*. Used in conjunction with phenotypic data on winter injury, establishment, and turf quality, this map allowed for the identification of seven winter injury associated QTL on chromosomes 8, 11, and 13. A BLAST study into these QTL of interest identified coding regions for proteins associated with abiotic stress tolerance in other species. Proteins of interest were associated with signal transduction pathways such as the mitogen-activated protein kinase pathway, as well as DRE family protein coding regions, both of which are linked to abiotic stress response and mitigation. These findings provided a compelling case for which proteins and pathways play a role in freeze tolerance.

Furthering our understanding of the physiological changes that zoysiagrass undergoes during cold acclimation may shed light on phenotypic traits useful in selection of more freeze tolerant genotypes. To investigate the relationship between cold acclimation, protein expression, and freeze tolerance in zoysiagrasses, *Z. japonica* cultivars with a reported differing level of freeze susceptibility and previously studied quantitative trait loci (QTL) associated with winter hardiness [24] were investigated here. The objectives of this study were to (i) compare the effects of cold acclimation treatments on freeze tolerance in these cultivars and (ii) to identify proteins associated with cold acclimation and freezing tolerance.

## MATERIALS AND METHODS

### Plant materials and growing conditions

Commercial cultivars ‘Meyer’ (Z. *japonica*, [25]) and ‘Victoria’ (Z. *japonica*, [26]) were selected for evaluation due to their reported range in freeze susceptibility (acclimated LT_50_ of −11.5°C and −9.5°C, respectively) [4, 5]. In the fall of 2015, cultivars were collected from research plots at the Lake Wheeler Turfgrass Field Laboratory (Raleigh, NC) and established in 24.5 cm × 50.8 cm plastic trays (Hummert International, Earth City, MO) containing Fafard P3 potting mix (Conrad Fafard Inc., Agawam, MA) in the greenhouse. In spring 2016, plants were vegetatively propagated into 2.5-cm-diameter, 12-cm-deep Ray Leach cone-tainers (Stuewe & Sons, Inc., Corvallis, OR) in USGA grade sand [27] using a single stolon or rhizome containing root, node, and shoot material according to Patton and Reicher [5]. Plants were allowed to establish for at least six weeks in the greenhouse at 27±2°C.

### Acclimation treatments

After establishment in the greenhouse, plants of each genotype were randomly arranged in a EGC walk-in chamber made by Environmental Growth Chambers (510 East Washington St. Chagrin, OH 44022) and cold acclimated (CA) for four weeks. The growth chamber used a consistent light emitting diode (LED) and temperatures of 8/2°C day/night cycles with a 10-h photoperiod of 300 µmol m^−2^s^−1^ photosynthetically active radiation [9]. Non acclimated (NA) plants of both genotypes remained in the greenhouse at 27°C during this four-week period.

### Evaluation of freezing tolerance

After acclimation treatments, intact plants in cone-tainers were evaluated for freeze tolerance based on whole plant survival. Immediately before placement in the freezer, NA plants were randomized with the CA plants. Based on the capacity of the freeze chambers, plants were arranged in each rack in a randomized complete block design with two replications. Each experimental unit consisted of 10 cone-tainers for each of the genotypes by acclimation treatment combinations. Based on reported LT_50_s [4, 5] and a pilot study in the spring of 2015, temperatures of −8°C, −9°C, −10°C, and −11°C were selected for this study. Each rack was placed in a plastic bag to minimize desiccation during freezing [11, 12] and placed into upright Kenmore refrigerators modified with cooling fans and FE Micro-controllers PXR4 (Price’s Scientific Services, Inc.). In the freezers, plants were initially held at −3°C for 15 h to remove latent heat from the soil. The temperature was then decreased at a rate of −1°C h^−1^ until reaching the target temperature, which was maintained for 3 h. Plants were thawed at a rate of 2°C h^−1^ until reaching the target temperature of 3°C. All trays were subsequently removed from plastic bags and placed in a walk-in growth chamber at 3°C for 24 h. Finally, plants were allowed to come to room temperature (22°C) overnight before being returned to the greenhouse. This experiment was repeated for a total of three runs. The second run of freeze tests was discarded because of a malfunction in the commercial freezers so only data from runs one and three were used in the analysis.

Plants were evaluated weekly for survival for four weeks after freezing. Survival data was taken on a binary scale of 0 (death) to 1 (survival) evaluating each individual plant for the presence of green tissue [15]. Survival data collected four weeks after freezing was evaluated with a logistic regression analysis. This method uses binary data to determine the probability of survival against the temperature gradient and estimate LT_50_s according to Dunne et al. [15]. The LOGISTIC procedure in SAS version 9.4 (SAS Institute; Cary, NC) with a stepwise selection method generates a model and maximum likelihood estimates that were used to calculate odds ratios, survival probabilities, and LT_50_s. Regression models were created to evaluate each cultivar by acclimation treatment combination as well as acclimation treatments with cultivars pooled. Temperature was treated as a continuous variable to account for variability in the commercial freezers. Non-acclimated Victoria was used as a reference variable for the generation of the logistic regression model of cultivar by acclimation treatment because it had the lowest reported LT_50_. In the model of acclimation treatments alone, the non-acclimated treatment was considered a reference variable.

### Proteomic analysis

After acclimation treatments, sixteen cone-tainers of each cultivar × treatment combination for each of three reps were randomly selected and destructively sampled for proteomic analysis. Meristematic tissues from the grass crowns was harvested with less than 2.5 cm of root and shoot tissue remaining, and bulked for each cultivar-treatment within each run. Fresh tissues were immediately frozen in liquid nitrogen, held at −80°C until completion of the experiment, and then lyophilized (Genesis Pilot, SP Scientific, Warminster, PA). Lyophilized tissues were then ground into a fine powder (Retsch Mill Mixer MM 400, Haan, Germany) to aide in protein extraction. Proteins were extracted from approximately 0.5 g lyophilized tissues using the trichloroacetic acid (TCA)/acetone/phenol method previously described by Xu et al. [28]. Total protein concentration was quantified according to the methods of Bradford [29] with bovine serum albumin as a standard. Separation of proteins based on first-dimension isoelectric focusing (IEF) was done by loading 300 ug of protein dissolved in rehydration buffer (8M Urea, 2% Chaps, 50mM dithiothreitol (DTT) 0.2% (w/v) 3/10 ampholytes, and trace amounts of Bromophenol Blue) onto immobilized pH gradient (IPG) strips (pH 3-10, linear gradient, 11 cm), which were then focused in a Protean i12 IEF Cell (Bio-Rad Laboratories, Hercules, CA, USA). Second-dimension sodium dodecyl sulphate-polyacrilamide gel electrophoresis (SDS-PAGE) was done on 12.5% Tris-HCL Precast Gels using a Criterion electrophoresis cell (Bio-Rad Laboratories, Hercules CA, USA), run at 80V for 20 min before switching to 100V for approximately 2 h. Gels containing protein spots were then stained using the ProteoSilver Plus Silver Stain Kit (Sigma-Aldrich, MO, USA) according to manufacturer instructions. Three technical replicates for each biological replicate were used for 2-DE, resulting in a total of nine gels per cultivar-treatment combination.

The 2-DE gels were scanned using a GE Typhoon FLA 9500 Phosphorimager (GE Healthcare, PA, USA). Scanned images were aligned and spot intensity was quantified using Melanie 8 (SIB Swiss Institute of Bioinformatics, Lausanne, Switzerland). Spot intensity was used to determine fold change between the non-acclimated and acclimated samples within each cultivar. Differences in protein expression between the cultivars and cold acclimation treatments were analyzed using ANOVA, and spots with a minimum of 2.0-fold difference in abundance (upwards or downwards) between cultivar, treatment, or interaction were selected. Spots that showed consistent changes throughout replicates were considered for spot excision, and selected spots were manually excised. The silver stained protein spots were de-stained using the ProteoSilver Plus Stain Kit (Sigma-Aldrich, MO, USA), and in-gel enzymatic digestion was done overnight at 37°C using Trypsin Singles, Proteomics Grade (Sigma-Aldrich, MO, USA) following manufacturer instructions for preparation of reagents.

The sample peptide solutions were analyzed using a MALDI-TOF/TOF mass spectrometer (Bruker UltrafleXtreme, Coventry, UK). The observed MS peaks obtained were compared using MASCOT. The NCBI non-redundant protein database using the Viridiplantae filter was used to obtain the most likely protein match based on a peptide mass fingerprinting, and only the top hits were used and a score threshold of 40 was used as a cut off. The search parameters were as follows: trypsin enzyme, fixed modification of cysteine as carbamidomethylated, peptide charge state of +1, peptide tolerance of 0.5 daltons, and up to 1 missed cleavage. For automatic, high throughput functional annotation, the closest Arabidopsis thaliana (AT) match was run through Blast 2 Go. For functional characterization, protein homologues were identified against the NCBI Arabidopsis database. Further functional information was gathered from the Arabidopsis Information Resource database (TAIR). Proteins were classified into categories based on their function [30].

## RESULTS AND DISCUSSION

### Freezing tests

Treatment, temperature and the interaction between them had significant type III effects based on the Wald *x*^2^ in the logistic regression model (Table 1). In the model that evaluated each cultivar by treatment combination, cultivar was not a significant effect (p = 0.6486) (Table 1a). However, when cultivar was excluded from the modes (Table 1b), acclimation treatment and temperature were both significant (p<0.0001).

**Table 1.**
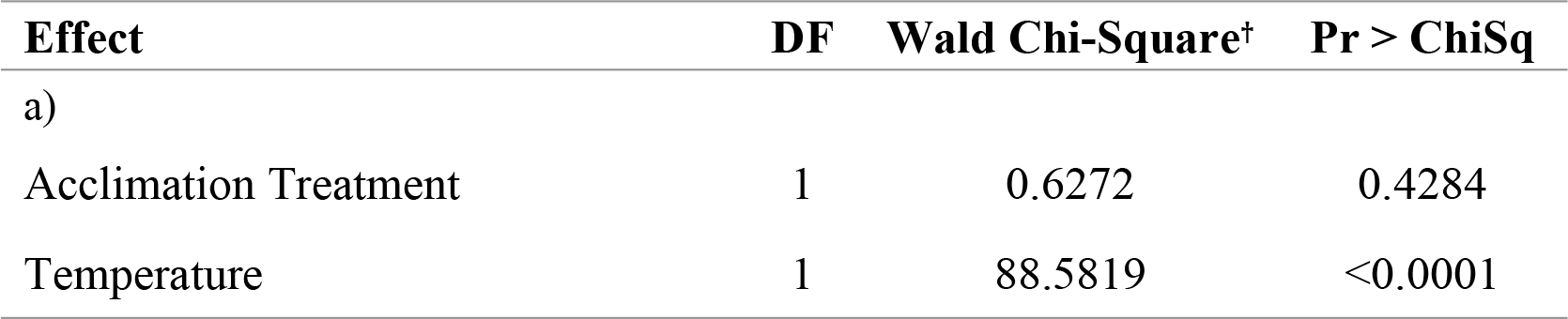
a) Type III analysis of stepwise-selection logistic regression model used to estimate LT_50_ values for freeze tests of zoysiagrass cultivars under controlled environmental conditions: a) with and b) without cultivar included in the model.

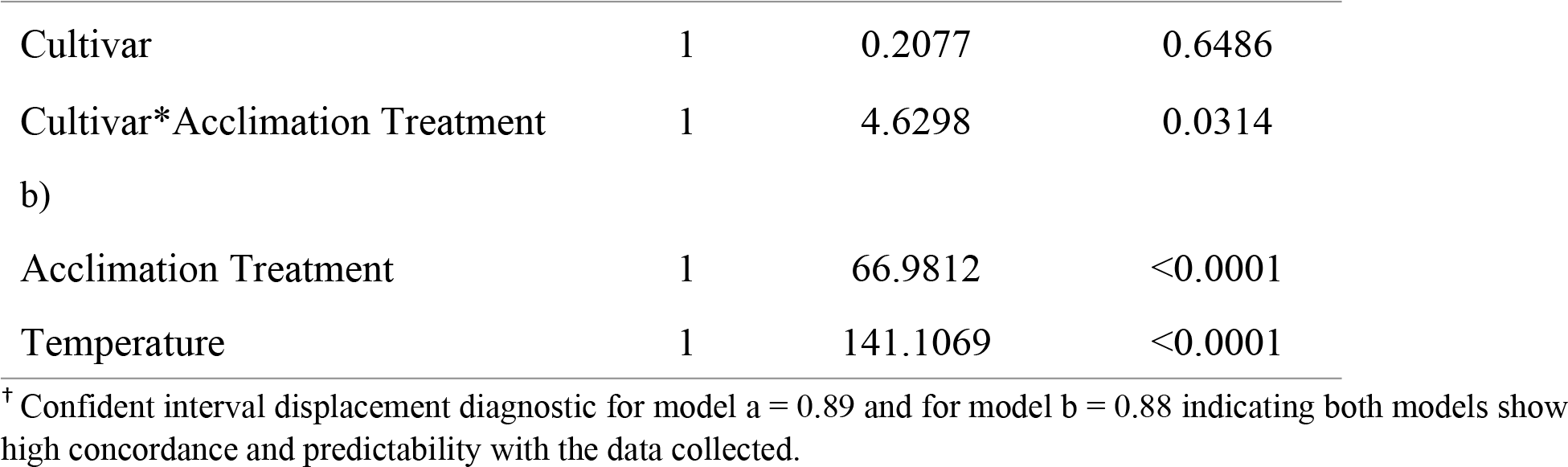

Maximum likelihood estimates generated from the logistic model were used to assess the survivability of each acclimation treatment or cultivar by acclimation treatment combination in relation to the reference variable by way of the LT_50_ (Table 2). Because NA Victoria was used as a reference variable, all estimates show differences in freezing tolerance in relation to it. The mean separation test indicated that cultivars within acclimation treatments were significantly different (p < 0.05). The cold acclimation treatment altered Meyer and Victoria’s LT_50_s by −2.66°C and − 1.50°C, respectively (Figure 1). Fitness of both these models showed high concordance and predictability with the data collected (Tables 1 and 2). Through modeling of acclimation treatments alone, CA plants were determined to be approximately 1.9-fold more likely to survive than NA plants (Table 2). Similar studies in zoysiagrass [4], St. Augustinegrass [11, 12, 13], bermudagrass [10], buffalograss [8] and saltgrass [7] have also reported that plants demonstrated better freeze tolerance when subjected to cold acclimation. Our results of a nearly two-fold increase supports these findings. Moreover, CA Meyer was significantly different from CA Victoria, but NA cultivars were similar to each other (Figure 1).

**Table 2.** Maximum likelihood estimates used to calculate survival probabilities and LT_50_ values in freeze tests of zoysiagrass under controlled environmental conditions: a) excluding cultivar from the model and b) for each cultivar by acclimation treatment combination.

| Parameter | Estimate<sup>†</sup> | Wald Chi-Square<sup>††</sup> | Pr > ChiSq |
| --- | --- | --- | --- |
| a) |  |  |  |
| Intercept | 7.6470 | 130.9546 | <0.0001 |
| Cold Acclimation Treatment | 1.8934 | 66.9812 | <0.0001 |
| Temperature | 0.8685 | 141.1069 | <0.0001 |
| b) |  |  |  |
| Intercept | 7.7784 | 124.5539 | <0.0001 |
| Meyer Acclimated | 2.6222 | 58.3716 | <0.0001 |
| Meyer Non-Acclimated | 0.1241 | 0.1860 | 0.6663 |
| Victoria Acclimated | 1.4277 | 22.1395 | <0.0001 |
| Temperature | 0.8902 | 140.8128 | <0.0001 |

<sup>†</sup> The reference variable used for the estimation of the logistic regression model was a) the non-acclimated treatment and b) Victoria Non-Acclimated. Temperature was treated as a continuous variable for both models.
<sup>††</sup> Confident interval displacement diagnostic for model a = 0.84 and for model b = 0.87 indicating both models show high concordance and predictability with the data collected.

**Figure 1.**
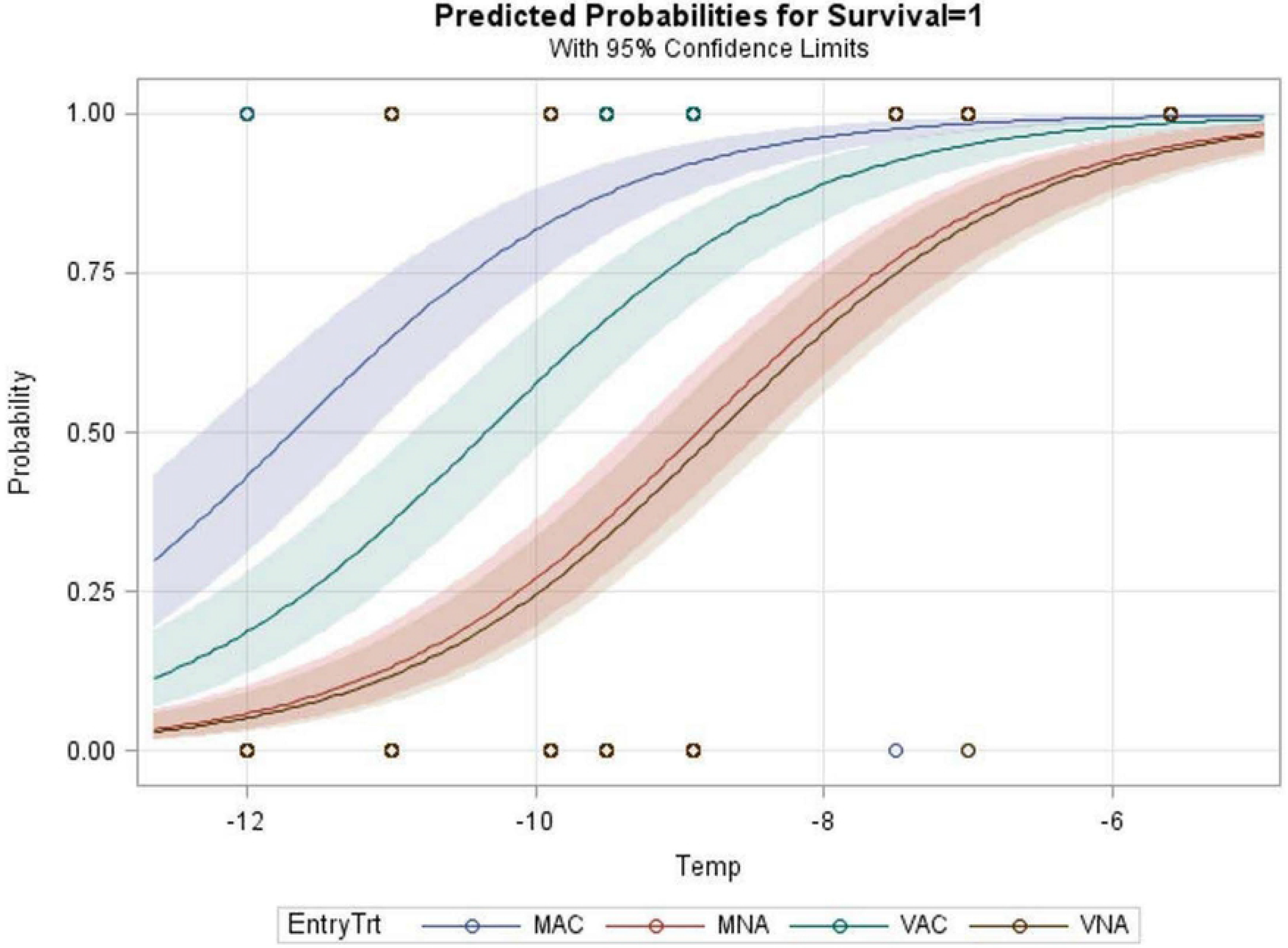
Probability of zoysiagrass survival in controlled environment freeze tests across temperature curves for each cultivar by acclimation treatment combination where MAC: acclimated ‘Meyer’ with an LT50 of −11.56°C, MNA: non-acclimated ‘Meyer’ with an LT50 of − 8.89°C, VAC: acclimated ‘Victoria’ with an LT50 of −10.30°C, and VNA: non-acclimated ‘Victoria’ with an LT50 of −8.75°C. LT50 values occur at probability 0.50.

Patton et al., [5] observed that specific carbohydrates, proline, and certain proteins in CA zoysiagrass were correlated with freeze tolerance in thirteen genotypes. A proteomic response to cold acclimation was also observed in velvet bentgrass (*Agrotosis canina*) [31] where CA plants were not only more freeze tolerant but showed thirteen different protein spots that were differentially expressed in CA versus NA plants. Further analysis of metabolic processes that occur during cold acclimation and how they differ between these cultivars, may explain differences in freeze tolerance between *Z. japonica* genotypes.

### Protein abundance

Variation in the abundance of specific proteins was assessed in response to acclimation treatment by way of 2-DE SDS-PAGE. The non-acclimated meristematic tissue samples of each cultivar were used as the baseline to control for variation between the treatments. A total of 471 protein spots were detected on the 2-DE gels. From those, 62 spots were statistically significant (p≤0.05) and exhibited a minimum of a two-fold change in response to cold acclimation for at least one cultivar (Table 3). There were significant differences in expression of these 62 proteins for Meyer and Victoria. In Meyer, 35 spots were higher in abundance in response to cold acclimation (12, 17, 37, 40, 44, 56, 61, 62, 64, 80, 90, 124, 162, 185, 187, 191, 201, 234, 243, 272, 293, 311, 321, 340, 348, 366, 371, 401, 403, 414, 424, 438, 443, 466, 468), while five spots were lower in abundance (86, 87, 223, 240, 261). In Victoria, 46 spots were found to be in lower abundance in response to cold acclimation (17, 40, 44, 61, 62, 80, 90, 104, 114, 116, 131, 142, 148, 162, 169, 184, 185, 187, 191, 193, 201, 216, 223, 238, 240, 243, 261, 265, 270, 285, 292, 293, 300, 310, 311, 317, 321, 340, 348, 366, 371, 375, 401, 424, 438, 467), and seven were in higher abundance (37, 59, 75, 86, 87, 403, 468) (Figure 2). When comparing the 31 protein spots identified in both Meyer and Victoria, 23 were found to increase during cold acclimation in Meyer but decrease in abundance in Victoria. Moreover, two proteins were significantly increased for Victoria, but decreased for Meyer in response to cold acclimation. Protein spots that were unique to each cultivar were identified in order to attempt to elucidate their role in cold acclimation.

**Figure 2.**
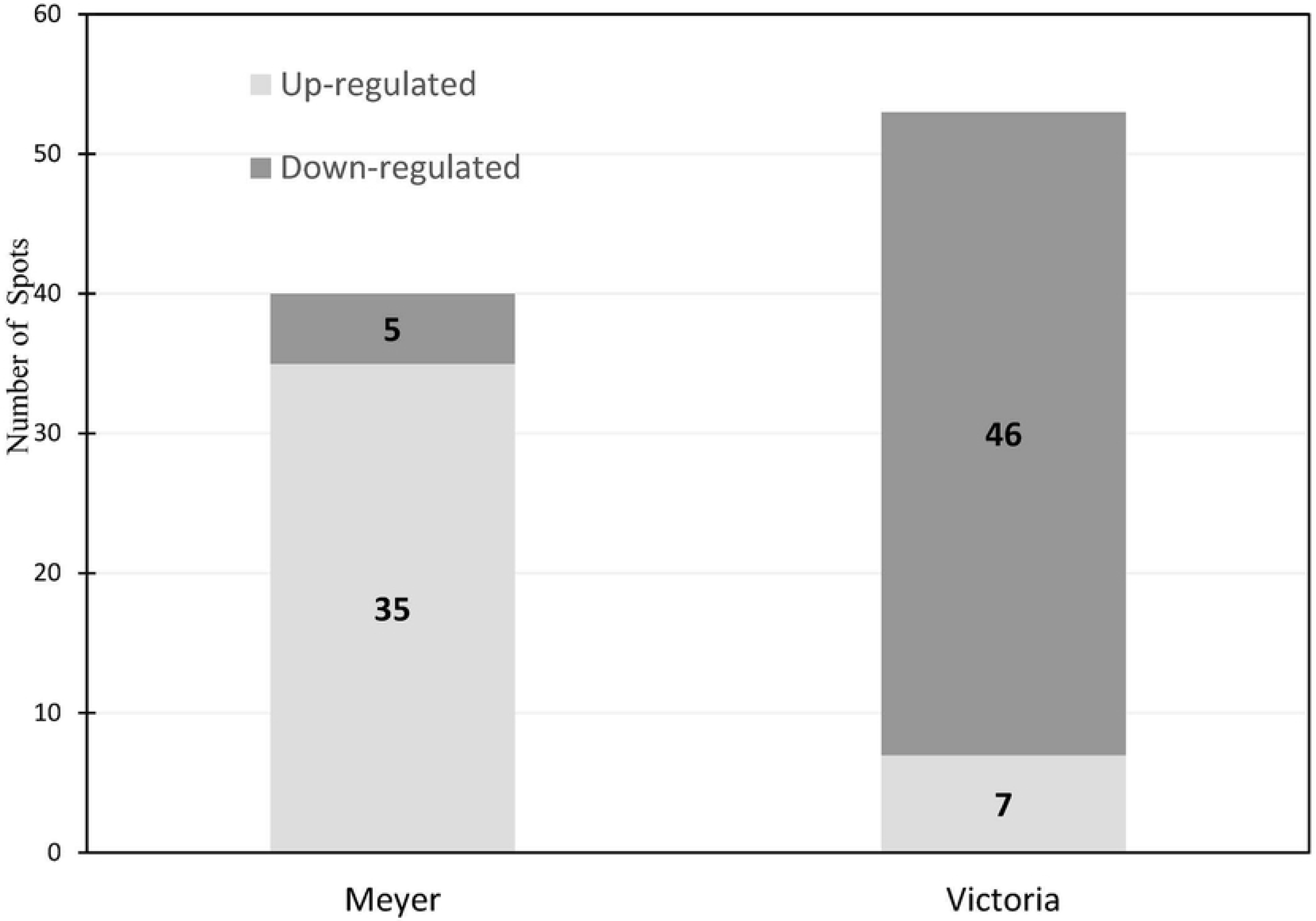
Number of protein spots identified in nodes of zoysiagrass cultivars ‘Meyer’ (freeze tolerant) and ‘Victoria’ (freeze susceptible) acclimated at 8/2°C (day/night) for four weeks, against a non-acclimated baseline.

In Meyer, eight unique spots were identified with at least a two-fold change in response to CA, all of which were upregulated (Table 3). These proteins were associated with late embryogenesis abundant protein 3, nudix hydrolase homolog 2, putative transmembrane protein (DUF506), manganese superoxide dismutase 1, general regulatory factor 2, Rossmann-fold NAD(P)(+)-binding protein, general regulatory factor 2, MAP kinase 10, and nucleotidyltransferase family protein. Among proteins that were upregulated in response to CA in Meyer, compared to the NA baseline, three proteins were identified as being of interest: Late embryogenesis abundant protein 3 (LEA3), with a 6.7 fold change during CA (one of the highest changes observed) has been found to be functionally associated with disease and defense, and serves as a chaperone protein, to protect proteins and membranes particularly in situations of dehydration such as drought and cold stress [32]. This is consistent with previous reports of increased production of dehydrins, soluble proteins in the LEA family, in freeze tolerant zoysiagrass cultivars [17, 18]. Second, manganese superoxide dismutase 1 (SOD), with an almost two fold increase, is another disease and defense related protein which assists with reducing oxidative stress, an important factor during cold acclimation when metabolism is slowing yet light levels are still high [33]. Third, MAP Kinase 10, or mitogen-activated protein kinase (MAPK/MPK), exhibited a three-fold increase. This protein plays a crucial role in signal transduction pathways for several biotic and abiotic stresses, and also serves to regulate C-repeat binding factor (CBF) gene expression during cold stress signaling in which the expression levels of CBFs determine the levels of cold-regulated (COR) gene expression and freezing tolerance [34, 35, 36].

**Table 3:**
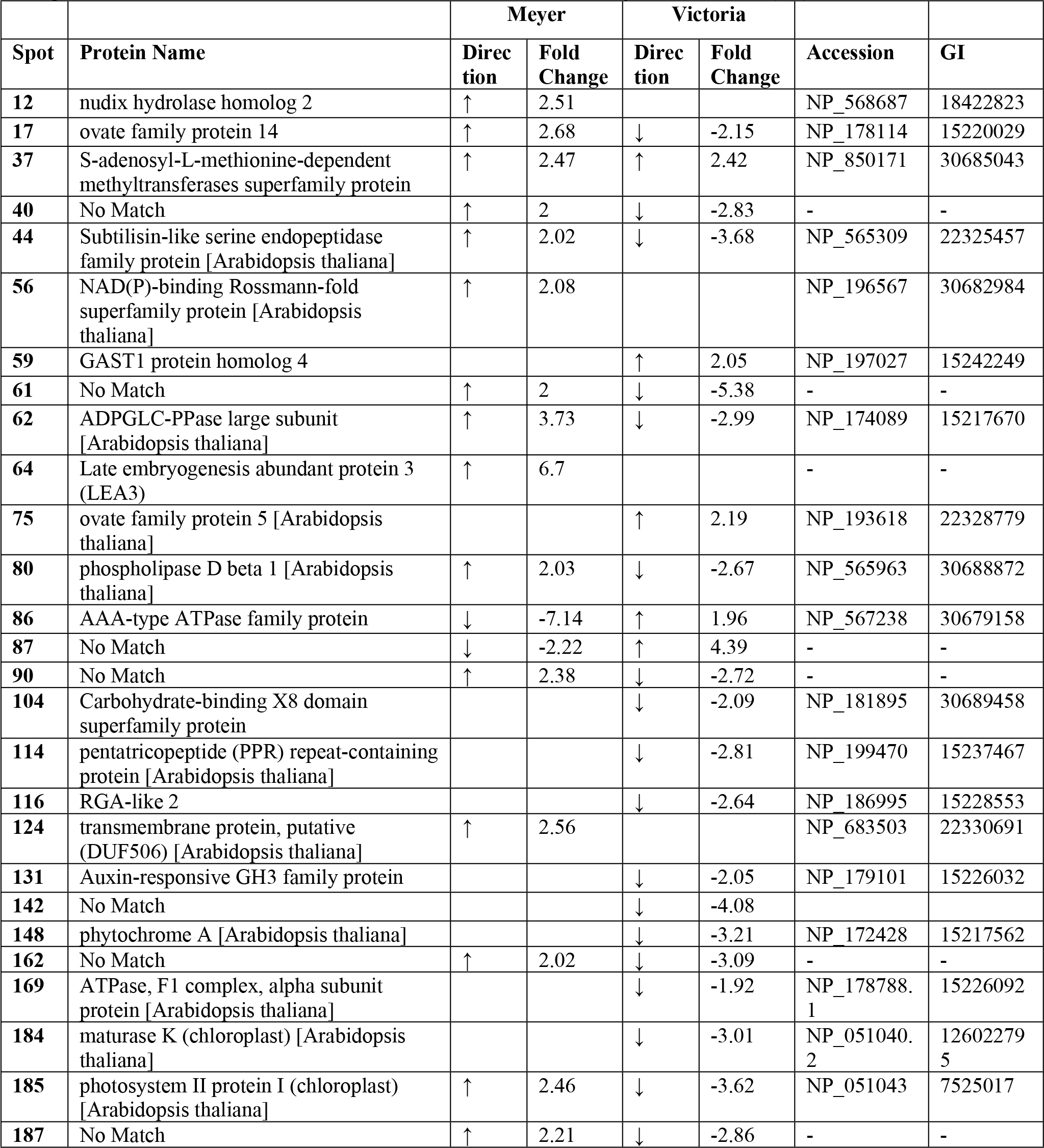
Functional data of the hypothetical proteins matching to each identified protein spot number as determined through Blast 2 Go. The protein name, fold change and direction of the change were determined and listed with the accession and Glycemic Index (GI) number.

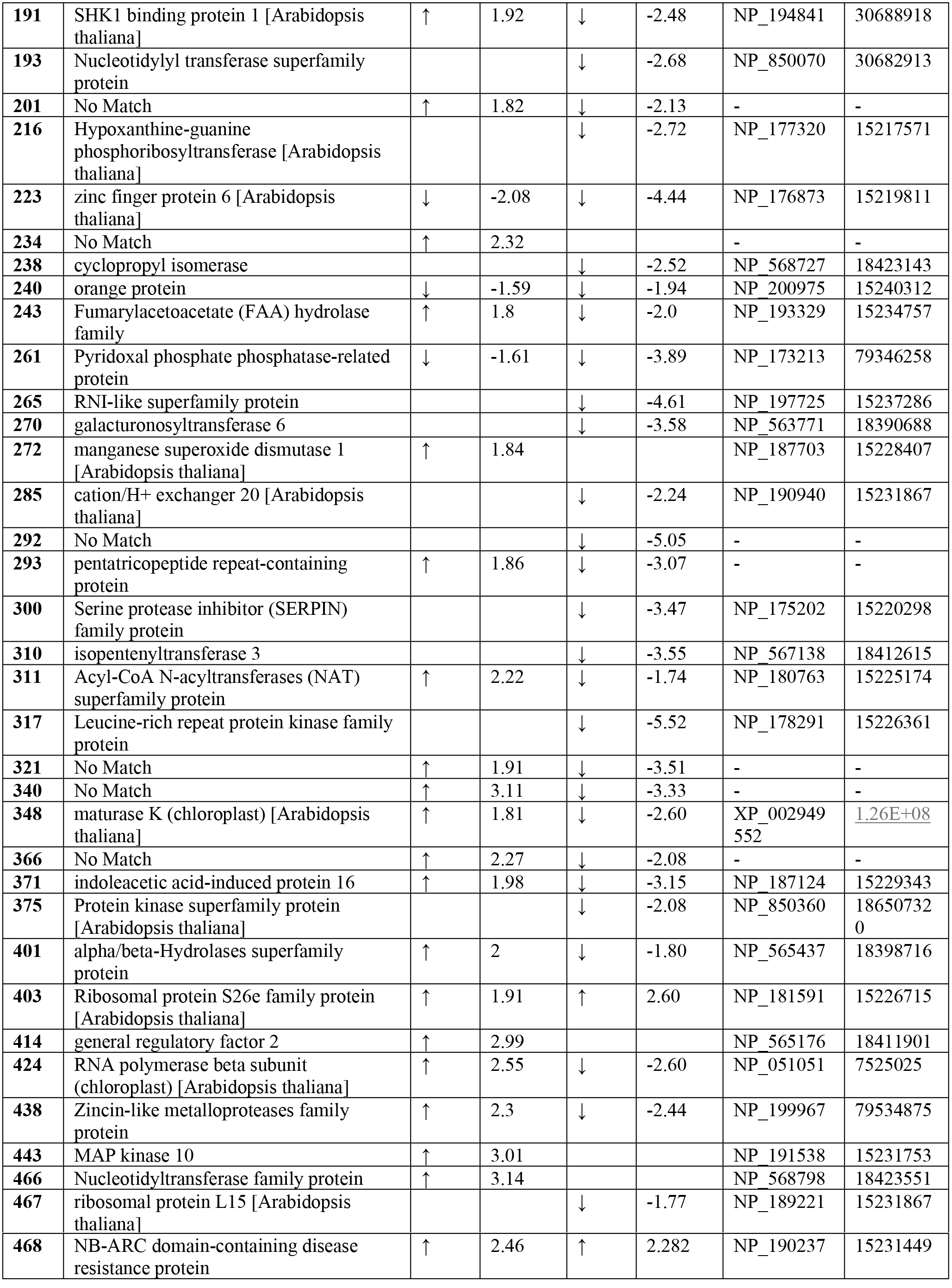

In Victoria, there were 20 identifiable, unique spots with at least a two-fold change, two of which were upregulated in response to CA, and 18 of which were downregulated (Table 3). Of these 20 spots, the seven proteins with the largest change in abundance were identified to be homologous with known proteins in online databases (Table 3). These predicted protein domains included Phytochrome A, GAST1 protein homolog 4, ovate family protein 5, leucine-rich repeat protein kinase family protein, galacturonosyltransferase 6, isopentenyltransferase 3, Maturase K (MatK), and RNA like superfamily protein. A few proteins of interest were identified through the differential expression in the NA vs CA treatments in Victoria. GAST1 protein homolog 4, an upregulated protein associated with cell growth and division, which serves as a GA-responsive protein [37]; Phytochrome A, with more than a 3-fold decrease, is a signal transduction related protein that plays a critical role in the cold response by sensing the changes in light conditions, and has been shown to promote *CBF* gene expression 9*CBF1*, a gene encoding AP2 domain-containing transcriptional activator that binds to the low-temperature-responsive element CCGAC and induces the expression of some cold-regulated genes, increasing plant freezing tolerance and subsequent cold tolerance [38]; ATPase F1 complex, alpha subunit protein, a down regulated energy related protein that extrudes protons from cells of plants to generate electrochemical proton gradients, which has a major role in providing the energy for secondary active transport across the plasma membrane, as well as playing a role in adaptation of plants to changing conditions, particularly stress [39].

Expression of the numerous shared proteins between Meyer and Victoria were determined to be of interest particularly if there was significant differential regulation of the proteins between the two cultivars as a response to CA. The shared cold responsive proteins of interest were: ADP-glucose pyrophosphorylase large subunit (upregulated in Meyer and downregulated in Victoria), a key metabolism related enzyme in the starch biosynthesis pathway [40]; Phospholipase D beta 1 (PLD) (upregulated in Meyer and downregulated in Victoria) which functions in metabolic processes, was implicated in the activation of cold response and may play an important, positive role in freezing tolerance [41, 42]; and Photosystem II protein I (PSII) (upregulated in Meyer and downregulated in Victoria), an energy related protein that could point to a better ability to keep photosynthesis active and therefore support energy needed for cold acclimation [43]. Glucose-1-phosphate adenylyltransferase large subunit (AGP), with a greater than three-fold upregulation in Meyer, and a greater than two-fold downregulation in Victoria, functions in the process of starch metabolism due to the significant role it plays in temperature induced yield loss, such as serving as a rate limiting step in starch biosynthesis under elevated temperatures in maize [44].

### Potential functions

Due to the relative lack of genetic information on turfgrasses, proteins of interest were identified based on homology with *Arabidopsis thaliana*, as well as through supporting data on their relation to abiotic stress tolerances associated with freezing. In the present study, 62 spots showed at least a two-fold change from the control treatment. The identified protein spots determined through mass spectrometry were sorted into categories/families according to their associated functions via the TAIR database. Out of the 62 identified protein spots, only 49 could be characterized by way of category/family and/or homologous protein. Proteins that were unnamed but classified were still included. However, proteins that could not be associated with a category/family were discarded from further analysis. Some overlap was present between the categories/families, but proteins were sorted according to their strongest match. Transcription and metabolism related proteins accounted for 31% and 21% of the total proteins, respectively. Energy related, and protein destination and storage related proteins made up 10% of the total identified proteins. Signal transduction related proteins consisted of four individuals (8%). The remaining categories of proteins were disease/defense and protein synthesis (3 proteins or 6% each), cell structure (two proteins or 4%), and cell growth/division and secondary metabolism (one protein or 2% each) (Figure 3).

**Figure 3.**
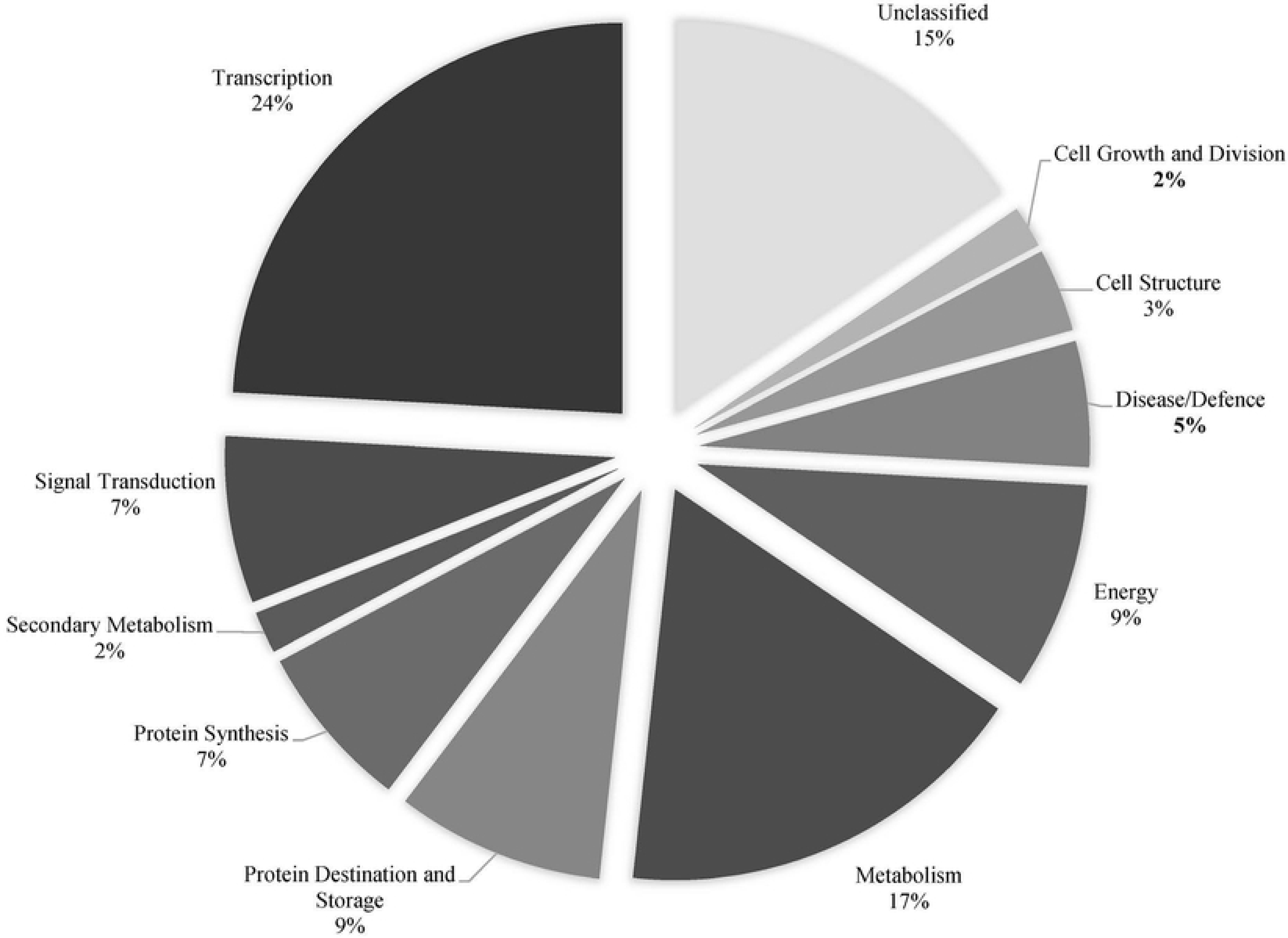
Functional categories (and their corresponding relative distributions) of proteins with a two-fold change or greater in response to cold acclimation.

#### Transcription

Transcription related proteins were primarily upregulated in Meyer and downregulated in Victoria, though there were a few exceptions. S-adenosyl-L-methionine-dependent methyltransferase super family was upregulated for both cultivars at a similar fold change. Zinc finger protein 6 was downregulated for both, but with a greater fold change for Victoria. The ovate family protein 5 transcriptional repressor was upregulated in Victoria. Todaka et al. [45] conducted a study in rice to investigate a large number of stress-responsive genes related to gene expression regulation, such as protein kinases and transcription factors, and direct plant protection against stress, such as metabolic enzymes, aquaporin, and late embryogenesis abundant proteins. Transcription factors play important roles in signaling cascades that lead to abiotic stress responses, by regulating the expression of downstream target genes via the utilization of cis-elements in the promoter regions of target genes under stress conditions. The dehydration-responsive element binding protein 1/C-repeat factor (DREB1/CBF) regulon has been identified in Arabidopsis to function in cold stress response, particularly in the accumulation of proline, regulation of stress-responsive gene expression, induction of transcription factor expression [46], and regulation of ABA responsive gene expression [47]. Overexpression of DREB1-type genes led to enhanced expression of stress inducible genes, including cold and drought stress, and elevated tolerance levels, an indicator that the DREB1 regulon may be one of the master regulatory systems for abiotic stress responses [45]. Therefore, a major fold change in transcription related proteins related to these regulatory systems might indicate a key effect response to cold stress.

#### Metabolism

Metabolism related proteins, such as phosopholipase D and glucose-1-phosphate adenylyltransferase were upregulated in CA Meyer, and downregulated in CA Victoria. Sugars are utilized by plants to counteract stressors in their natural environment. As a response to evolutionary pressures in stressful environments, there was a diversification of sugar structures and functions, most importantly the ability to modulate the expression of genes related to abiotic and biotic stress response systems through sugar/starch metabolism. Soluble sugars, as well as amino acids, polyamines, and polyols are all known to contribute to the development of cold stress tolerance mechanisms [48]. Saccharides can stabilize phospholipid membrane composition to maintain fluidity under cold stress conditions [49]. Nitrogen reserves in the form of increased abundance of certain amino acids, may function to improve overwintering and regrowth ability in plants [50]. The differential regulation of glucose-1-phosphate adenylyltransferase, supports the importance of starch metabolism. In addition to being the main storage carbohydrate in plants, starch provides a steady supply of energy when photosynthesis is not possible, resists water loss in stomatal pores, and aids cell differentiation. Under abiotic stresses, starches are remobilized to provide energy when photosynthesis becomes limited. Fructan, an important storage carbohydrates, is known to stabilize cell membranes to prevent water leakage during freezing [51]. Other sugars and metabolite derivatives support growth under stress conditions, as well as mitigate some of the negative effects of the stress by activating stress response cascades with signaling sugars [52].

#### Protein destination and storage

Protein destination and storage related proteins were upregulated in Meyer and downregulated in Victoria. Serpins play a critical role in the control of proteolysis, through inhibition of proteinases. Proteolysis is in turn crucial for plant stress response, defense, growth, and development. Silverman et al. [53] purports that cell lysis as a result of cold stress may release proteinases and damage cells. Therefore, plant serpins may act in a manner similar to mammalian serpins to protect cells from proteinases. The Shk1 kinase binding protein 1 (SKB1) is a floral initiator that responds to salt stress by suppressing transcription of flowering locus c (FLC) and several stress responsive genes [54]. FLC is also modulated by a flowering autonomous pathway gene (FLV) when cold temperatures are sensed to initiate cold response [55]. The upregulation of destination and storage related proteins in freeze-tolerant Meyer may serve to mediate and increase response time to damage caused by abiotic stress.

#### Energy

Proteins related to energy were differentially expressed between the two cultivars under the different acclimation treatments. Pyridoxal phosphate phosphatase-related protein was downregulated for both Meyer and Victoria CA cultivars compared to non-acclimated, with a greater fold change for Victoria. Meanwhile, ATP synthase had a two-fold downward change in Victoria only. The AAA-type ATPase family protein had a greater than two-fold downward change in Meyer, and a two-fold upward change in Victoria. Conversely, photosystem II protein I (chloroplast) had a greater than two-fold upward change in Meyer, and a greater than two-fold downward change in Victoria. Proteins related to photosynthesis, such as ATPases, are sensitive to changes in the environment, particularly those which result in a change in photoperiod, such as the changing of seasons into winter. This sensitivity is due to the need to balance light energy absorption with energy consumed by plant metabolism during the process of photosynthesis. Additionally, low temperatures have been found to heighten the disparity between light absorption and metabolic needs, making it necessary to adjust photosynthesis systems to maintain balanced energy flow. There is supporting evidence that the processes which interact with photosynthesis, particularly during cold acclimation, involve several pathways leading to the regulation of acclimation to cold temperatures, including photosynthetic redox, acclimation, and sugar signaling pathways [56].

#### Protein synthesis

Proteins related to protein synthesis were upregulated in Meyer, and primarily downregulated in Victoria, with the exception of spot 403: ribosomal protein S26e family protein, which was upregulated in Victoria. The decrease in ribosomal proteins overall may play a role in translation, ribosome assembly, and proper function of mRNA under low-temperature conditions [57]. A study in tall fescue [*Schedonorus arundinaceus* (Schreb.) Dumort. syn. *Festuca arundinacea* Schreb.] demonstrated that the introduction of the isopentenyltransferase (IPT) gene enhanced cold tolerance, as well as tillering ability and chlorophyll a and b levels [58]. This gene is purportedly responsible for inhibited apical dominance, delayed senescence, and increased chlorophyll and secondary metabolite levels [59]. Research in peanut (*Arachis hypogaea* L.) [60] and sugarcane (*Saccharum* spp. L.) [61] support the ability of IPT to delay stress induced plant senescence, resulting in enhanced drought and cold tolerance. Synthesis related proteins such as IPT likely play an important role in generating cold acclimation induced proteins.

#### Signal transduction

Signal transduction related proteins were upregulated in Meyer, but downregulated in Victoria. Additionally, a higher number of significant changes in signal transduction related proteins occurred in Victoria at a 3:1 ratio. Research in *Arabidopsis thaliana* identified calcium as the most common signaling factor for cells to translate external stimuli into a biochemical response. In the presence of cold stress, calcium levels are elevated, and are correlated to the activation of certain chilling sensitive genes [62]. A decrease in endogenous levels of IAA-amido synthetase, as observed in Victoria, was shown to enhance drought tolerance in rice in conjugation with TLD1, as well as influence an increase in LEA gene expression [63]. A decrease in free IAA led to enhanced ROS scavenging and increased cold response and membrane permeability as well, leading to increased cold tolerance [64]. MAP kinase 10, the highly upregulated protein observed in Meyer, is known in several species, including *Arabidopsis thaliana*, tobacco (*Nicotiana tabacum*), tomato (*Solanum lycopersicum*), and rice (*Oryza sativa*), as a major player in plant stress signaling. The MAPK cascade is a major pathway for multiple plant stress responses through its involvement in mediating oxidative stress, as well as its interaction with several other important signaling pathways [65]. Signal transduction related proteins may be of particular interest in breeding due to their key roles in stress response pathways.

#### Disease and defense

While a small percentage of proteins related to disease and defense showed a significant fold change, all identifiable proteins showed two-fold or greater upregulation. The two proteins of particular interest in Meyer were LEA3 and SOD. Kobayashi et al., 2004 observed that LEA proteins are a major downstream group involved in the ABA-dependent and independent signaling pathways for freezing tolerance in wheat and may be able to be further enhanced for increased response. Genes in the LEA family commonly encode highly hydrophilic proteins, which positively correlate with greater freezing tolerance when overexpressed. Additionally, in a similar manner to cold acclimation, light conditions influenced the accumulation of cold responsive LEA proteins. Transcriptional processes were enhanced by light and suppressed by darkness in the Kobayashi et al. [66] study. SODs operate as a first line of defense against reactive oxygen species, which are increased under stress conditions. Exposure to oxidative stressors results in an increase in SODs, and induced overexpression of SODs has led to increased protection against specific stresses, though results have been mixed due to the complexity of an associated scavenging pathway [33]. However, some attempts to overexpress SOD have been successful in increasing stress response [67], and may be utilized in future breeding efforts.

#### Other proteins

Cell structure, cell growth and division, and secondary metabolism related proteins made up the smallest percentage of the significant fold changes between the two cultivars. The plasma membrane is the primary site of freezing injury, and maintaining an intact membrane can act as an effective barrier to the formation of ice crystals [68]. The 2.5 upward fold change of a transmembrane cell structure related protein in Meyer may indicate a role in the maintenance of this membrane for protection against freezing injury. GAST1 protein homolog, associated with cell growth and division, is involved in numerous biological processes, including elongation, flowering, seed development and light signaling, flowering and stem growth, shoot elongation and flower transition. Rubinovich and Weiss [37] proposed that any sort of higher growth/elongation during the cold acclimation period could potentially interfere with proper cold acclimation, such as using photosynthates towards growth rather than protection. Little information exists on the pentatricopeptide repeat contain protein, other than it may be a product of partial degradation [69].

In comparison to Xuan et al. [23], their study found more differential expression between cold acclimated Meyer (*Zoysia japonica*) compared to Diamond (*Zoysia matrella*), whereas our study indicated a greater change in abundance in Victoria compared to Meyer. The use of cultivars of different species in their study might provide some explanation into this difference in results. Some of the differential expression observed in that study might be attributed to inherent differences between the species rather than actual differences in freeze response. They found that energy and metabolism were the largest functional categories at 23% and 21% respectively, as well as photosynthesis (13%), signal transduction (13%), and redox homeostasis and defense response (14%). While we also found these categories to be important, we had 31% of identified proteins from meristematic crown tissue as falling into the transcriptional category, in comparison with the 2% of stolon proteins associated with transcription by Xuan et al. The amount of transcription related activity may be significantly higher in crown tissue compared to stolon tissue, and therefore may be more involved in meristematic tissue growth compared to stolon growth. There was some overlap in identified proteins, particularly MAP kinase, Photosystem II, and phosphatase related proteins. Although there were several different proteins and pathways found through proteomic analysis, their study came to a similar conclusion that signal transduction and photosynthesis may explain some of the differences in freeze tolerance between Meyer and less freeze tolerant cultivars such as Diamond or Victoria. ADP glucose pyrophosphorylase (ADPGLC-PPase) and other similar ATPases were identified as potentially playing an important role in carbohydrate metabolism under cold stress by Xuan et al. [23], and was also observed in our study to be significantly upregulated in Meyer and downregulated in Victoria.

The study by Holloway et al. [24] identified 146 proteins that were potentially associated with abiotic stress response related pathways, a number which corresponded to the putative winter hardiness QTL that had been identified through SNP-based linkage mapping. Some overlap between those previously identified winter injury stress associated proteins and the cold acclimation associated proteins in this study were found. Leucine-rich repeat receptor-like protein kinase and pentatricopeptide repeat family protein are among the most notable corresponding proteins, as well as several related kinases and auxin related proteins. The presence of these proteins in both studies strengthens their association with cold stress response. Further analysis will be of great importance to confirming the findings of relevant cold acclimation associated proteins in different tissues of *Zoysia* spp.

### Overall cultivar differences

The uniformity in freeze tolerance of NA cultivars suggests that the physiological changes that occur during cold acclimation may be the separating factor between freeze tolerances of zoysiagrass cultivars. The large disparity in response to cold acclimation in Meyer and Victoria is of particular interest requiring further study. The large number of down-regulated proteins observed in Victoria in response to cold acclimation might indicate a differential way in which freeze tolerant and freeze sensitive cultivars physiologically respond to cold stimulus. The upregulation of proteins in freeze tolerant Meyer may drive the acclimation process, while the downregulation of proteins in freeze sensitive Victoria may be the result of shutting down processes to conserve energy. Victoria was originally selected by its developer for its ability to perform well under California growing conditions of drought and high temperatures with the ability to remain green during the onset of winter dormancy [26]. Conversely, Meyer zoysiagrass originated from Korea and was selected in Virginia and Maryland for its survival in the Northeast and Midwest despite its “disadvantage” of poor winter color [25, 1]. Under stress conditions, reduced photosynthesis can limit carbon supply and biomass production, resulting in reduced growth and fitness of the plant. However, plants that utilize starch remobilization can avoid these dangers by providing an alternate source of energy and carbon to survive and even mitigate the effects of a stressful environment [52]. The high level of downregulation in unique protein spots in Victoria, may be a contributing factor of reducing photosynthesis, and therefore sensitivity to temperatures possibly allowing for improved fall color but reduced cold acclimation. Meyer adapts to cold temperatures by upregulating protein expression, and has shown higher levels of starch metabolizing proteins that may help with remobilization. However, further investigation is needed to verify these hypotheses.

Freeze tolerance is believed to be correlated with the accumulation of soluble proteins, notably by a general increase in winter and subsequent decline in spring [68]. However, as observed in this study, it is not always an increase in protein accumulation during cold acclimation, but rather about overall quantitative changes that occur during cold acclimation which contribute to freezing tolerance. Although the largest percentage of identified proteins were functionally related to transcription and metabolism, the relative importance of these proteins to cold acclimation and freezing tolerance is still being researched. Based on the functional data behind homologous proteins in related species, the major proteins of interest for their association with cold acclimation are LEA 3, MAPK, SOD, GAST1, Phytochrome A, ATP synthase, AGP, PLD, and PSII. The diversity in the metabolic pathways these proteins follow further emphasizes the complexity of abiotic stress response systems. The differential reactions between Meyer and Victoria to cold acclimation highlight the need for greater depth of knowledge on the mechanisms behind cold and freeze stress response, particularly the metabolic differences. Determining the nature of protein composition changes in the crown tissue of zoysiagrass genotypes following cold acclimation will aid in the development of new breeding strategies to produce cultivars with increased freezing tolerance.

## Acknowledgements

This research was supported in part with funding provided by the NC State University Plant Breeding Consortium, the NC State University Center for Turfgrass Environmental Research and Education, and the United States Golf Association.

